# A biomathematical approaches models to identify human platelet activation signature in response to various agonists

**DOI:** 10.1101/2025.02.06.636809

**Authors:** Fabrice Cognasse, Kim Anh Nguyen, Marco Heestermans, Charles-Antoine Arthaud, Marie-Ange Eyraud, Amelie Prier, Simon de Bernard, Julien Nourikyan, Anne-Claire Duchez, Stephane Avril, Olivier Garraud, Hind Hamzeh-Cognasse

## Abstract

**Background:** Platelets are crucial mediators at the crossroads of hemostasis, immunity, and inflammation, adapting their responses to diverse stimuli. Despite their recognized role, the precise pathways and markers associated with platelet activation remain poorly understood. This study aimed to unravel these mechanisms by evaluating platelet responses to various agonists, employing biomathematical models to map activation patterns and identify key biomarkers.

**Methods:** Using samples from ten healthy donors, platelets were exposed to seven stimulation conditions: unstimulated, PAR-1 agonist TRAP, PAR-4 agonist AYPGKF, ADP, collagen, sCD40L, and fibrinogen. A comprehensive analysis of 47 biological markers—covering membrane activation, soluble mediators, and signaling pathways—was conducted. Statistical and machine learning models, including hierarchical clustering and random forests, were applied to classify and interpret platelet activation signatures.

**Results:** Distinct activation profiles were observed for each agonist. A streamlined panel of six markers—AKT, CD40 ligand, CD62P (mean fluorescence intensity and percentage), PKC, RANTES, and TSLP—achieved 86.8% accuracy in identifying the activating stimulus. The study highlighted significant variations, influenced by both the stimulus and donor-specific factors. Machine learning approaches further refined classification, achieving a multiclass accuracy of 87.9%. Hierarchical clustering demonstrated clear distinctions, particularly between PAR-1/PAR-4 responses and other agonists.

**Conclusion:** This innovative research redefines platelets as dynamic "biological sensors" capable of decoding complex danger signals. By integrating biomathematical modeling and artificial intelligence, it identifies a precise biomarker panel with transformative potential for diagnostics and therapies in inflammation and immune disorders. This work positions platelets not just as key players in hemostasis but as programmable agents for precision medicine, heralding a new era in adaptive, AI-driven healthcare solutions.

**Author Summary:** Platelets are often seen as simple players in blood clotting, but they do much more. They sit at the crossroads of hemostasis (stopping bleeding), innate immunity (our body’s first defense), and inflammation. They even influence adaptive immunity and play key roles in maintaining healthy blood vessels and contributing to disease.

What makes platelets fascinating is their ability to respond quickly to their environment. They carry various receptors and release substances like growth factors, immune signals, clotting factors, RNA, and tiny vesicles. This helps them react to threats and communicate with other cells. But the big question is: can platelets tailor their response based on specific stimuli?

In my research, I set out to answer this. Using mathematical models and analytical techniques, I studied how platelets react to different triggers, especially those linked to immune and clotting responses. My goal was to identify specific molecular "signatures" that define how platelets respond.

I found that platelets can distinguish between danger signals and adjust their secretory responses. Normally, this helps manage threats efficiently. However, when this response exceeds what’s needed, it can contribute to diseases like cardiovascular disorders, severe infections, autoimmune conditions, and cancer.

Understanding these pathways opens new doors for treatment. Since platelet activity can be influenced by drugs, we could shift their role from harmful to beneficial in many diseases. This could revolutionize how we approach conditions driven by inflammation and immune dysregulation.

**Graphical Abstract:** 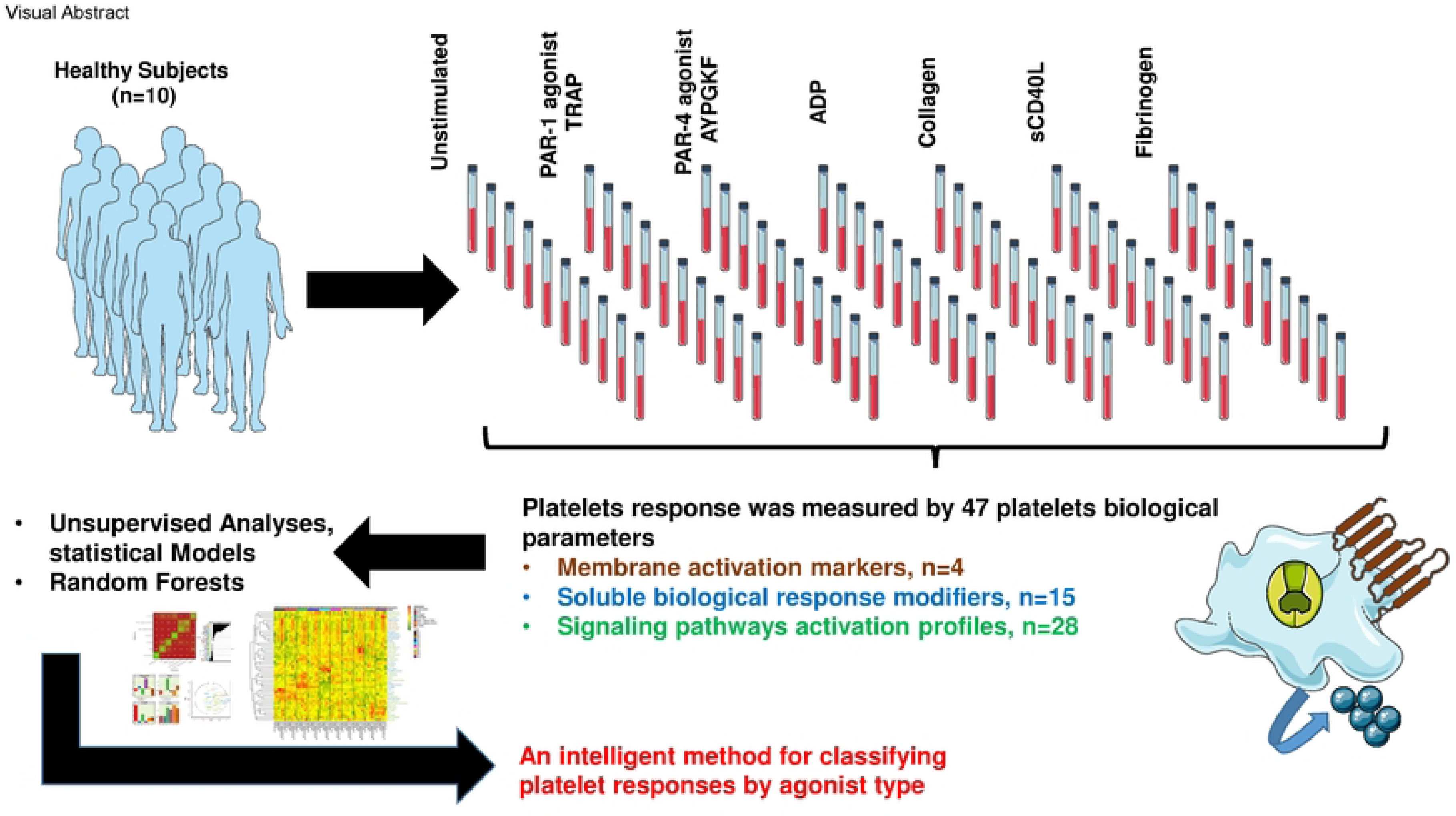

## Introduction

Platelets function at the intersection of hemostasis, innate immunity, and inflammation; moreover, platelets may also influence adaptive immunity and orchestrate vascular physiology and pathology. Platelets express numerous receptors and contain several secretory products that are instrumental in functional immune responses; thus, platelets provide potential new routes for drug targeting in inflammatory disorders [1–5]. They have been shown to play dedicated roles in innate immunity and pathogen sensing. Platelets express Toll-like receptor (TLR) family proteins on their surfaces [3, 6–8]. These platelet TLRs recognize pathogen-associated molecular patterns (PAMPs) and damage-associated molecular patterns (DAMPs) [9, 10], which are non-self-molecules and molecules associated with pathogen infection, respectively. In response to this stimulation, platelets deliver differential signals for the secretion of cytokines and chemokines.

Platelets are commonly accepted as secretors of proinflammatory cytokines, such as soluble CD40 ligand (sCD40L)/soluble CD154, chemokines and biological reaction modifiers, also known as immunomodulators [3, 11–14]. The related intracellular signaling mechanisms are less evident in platelets than in nucleated cells, the role and function of transcription factors remain to be explored. Notably, Lannan KL *et al.* [15] discovered the presence and nongenomic roles of transcription factors such as nuclear factor kappa β (NFκB) family proteins and peroxisome proliferator-activated receptor gamma (PPARγ) in platelets. Platelet-derived microparticles (PMPs), the small plasma membrane vesicles (0.05-1µm) released from platelets when they are activated, are the largest pool of microparticles circulating in the blood and are considered to be mediators of intercellular communication that trigger responses including inflammation, angiogenesis and tissue regeneration. Moreover, platelet microparticles convey an elaborate set of transcription factors and nucleic acids and contain mitochondria. Duchez AC *et al.* [16, 17] observed that microparticles and their cargo are internalized by activated neutrophils in the endomembrane system via 12(S)-HETE.

Platelets are highly responsive and secretory cells. They can alter their environment by releasing growth factors, chemokines, coagulation factors, RNA species and extracellular vesicles. On the other hand, platelets will also adapt to their environment. Platelets are able to sense danger signals and adapt their response depending on the stimulus; for example, platelets exhibit this behavior in an infectious context [18]. Platelets appear to be able to discriminate between different isoforms of lipopolysaccharide (LPS) that differ slightly in structure [19]. This outcome was confirmed by Kappelmayer *et al.* [20], who observed that the rough form of *S. minnesota* LPS (Re-LPS) was more likely to enhance platelet activation than smooth *E. coli* LPS. In addition, different LPSs from *enterohemorrhagic E. coli* serotypes showed different abilities to bind to platelets compared to the *nonenterohemorrhagic* serotypes at a similar dose, confirming the ability of platelet TLR4 to discriminate between closely related molecules [21]. It has been suggested that α-granules may be divided into distinct subpopulations and released by a stochastic process or, conversely, that they are differentially regulated in the release of angiogenic mediators in response to platelet agonists [22]. Despite this controversy, it is well known that activated platelets modulate angiogenesis by the differential release of pro- and anti-angiogenic factors from platelet α-granules in an agonist-dependent manner [23–25]. Using biomathematical approaches to assess and model platelet activation stimulated by different agonists, the role of activated platelets in angiogenesis could be deciphered.

Moreover, platelet activation contributes to signalosome mobilization due to changes in the vascular endothelium or invading infectious pathogens. Although several groups have convincingly demonstrated the links between the apparently distinct pathways of coagulation and inflammation, which are both complex responses to danger signals [26, 27], the clinical relevance of this interaction remains unclear.

In the current study, our objective was to decipher the pathways utilized by these cells to respond specifically to these stimuli. We applied a combined analytical and mathematical approach to investigate whether different platelet agonists, functioning via hemostatic or immune receptors, induce a platelet differential response. After platelet stimulation using six different agonists, we measured 47 platelet biological parameters (membrane activation markers, soluble biological response modifiers, and signaling pathway activation profiles).

Unsupervised analyses, statistical models and random forest models were applied to explore the resulting data and to present an unbiased method for classifying platelet responses by agonist type. Our most important data showed that a donor effect was responsible for a large part of the variability that was observed. The response of some variables was a function of the specific stimulus, whereas the response of others was a function of the donor or even the sample. Finally, we observed that AKT, CD40 ligand, CD62P (MFI and %), PKC, RANTES and TSLP, with an accuracy of 86.8%, were the best (limited) panel for the measurement of human platelet activity.

## Results

Platelet-rich plasma (PRP) was incubated with (or without) the PAR-1 agonist TRAP, PAR-4 agonist AYPGKF, ADP, fibrinogen, collagen or sCD40L; All these agonists are well known. Fibrinogen in the PRP frequently leads to uncontrollable coagulation and platelet activation, we also wish to add fibrinogen to our stimulation panels to observe possible modulation of platelet activation markers in the presence of a significant amount of fibrinogen. They are classically used in the study of platelet physiology and all these platelet signaling events converge upon the final common pathway of platelet activation [28]. Then, 47 biological parameters of platelets (4 membrane activation markers, 15 soluble biological response modifiers, and 28 signaling pathway activation profiles; hereafter collectively referenced as “variables”) were measured (**Supplemental Figure 1 and Supplemental Table 1)**; the data were then subjected to further analyses. We depict log-likelihood profiles for a variety of platelet activation markers after transformation using the Box-Cox method. Each plot represents the behavior of a specific marker under varying transformation parameters (λλ), which were applied to improve data normalization and enhance interpretability during statistical modeling. Some markers (e.g., CD62P-MFI, AKT) show a steep and well-defined peak, suggesting a strong response to a specific transformation. Others, like TSLP, appear flatter, implying a more uniform response to different transformations, likely reflecting underlying data characteristics. The variation in λλ values across markers highlights the heterogeneity of platelet activation profiles and their dependency on specific agonists or donor characteristics (**Supplemental Figure 1).** Markers such as AKT, CD62P (MFI and %), PKC, RANTES, TSLP, and others show high significance under specific stimulations. Some markers are broadly responsive (e.g., CD62P across multiple stimuli), while others are specific to a single condition (e.g., TSLP for PAR-4). Different agonists (PAR-1, PAR-4, ADP, etc.) elicit distinct activation signatures, PAR-4 induces strong responses in SYK, TSLP, and others and Collagen activates markers like RANTES and sCD62P. Some markers (e.g., CXCL9) show variation attributed to donor-specific factors rather than the stimulus **(Supplemental Table 1).**

To ensure suitability for unsupervised analyses, any data demonstrated to be nonnormally distributed was transformed; logit transformation was used to make the data approximately normal before performing the regression analysis, and log10 transformation was used to increase legibility by correcting for large differences in abundance between samples. To determine an appropriate transformation for the remaining variables, the log-likelihoods of the “Value ∼ Stimulation + Donor” model for various values of the 𝜆 parameter of the Box-Cox power transformation were computed. **Supplemental Figure 2** shows the distribution of the variables after transformation for the 10 donors, and summary statistics are described in **Supplemental Table 2**.

Each marker demonstrates a unique distribution pattern, reflecting its response to different stimuli. For example, markers such as CD62P-MFI or TSLP exhibit distinct peaks or wider variability depending on the stimulation. The unstimulated condition often shows a narrower or baseline distribution, indicating low marker expression in resting platelets. PAR-1 and PAR-4 agonists show broader or elevated distributions for specific markers, indicative of strong activation. Log-transformed scales are used for markers like AKT or CD62P-MFI, improving visibility of marker dynamics across a wide range. Some markers show symmetrical distributions, while others are skewed, reflecting donor-specific variability or differences in marker response sensitivity. Markers like CXCL9 and RANTES show overlapping distributions under certain stimuli, suggesting shared or linked pathways of activation. Others, like TSLP, reveal more distinct shifts, emphasizing their role in specific activation pathways. These plots validate that platelet activation is stimulus-specific and depends on the biological marker measured. The figure supports the study’s conclusion that a select panel of markers can reliably differentiate agonist-induced platelet responses (**Supplemental Figure 2).**

Pairwise comparison of the centered and scaled variable values after transformation was performed. For each pair of variables, the values were compared in all the samples (top right), and the corresponding correlation coefficient was reported (bottom left) (**Supplemental Figure 3)**. Perfectly correlated variables would draw a diagonal line and would have a correlation coefficient of 1 (bright green square). While some variables are strongly correlated, most are not. As expected, a very good correlation was observed between the expression of the platelet activating receptors CD62P and CD63 (e.g. CD62P−MFI (log10) vs CD63−MFI (log10): 0.928) and between the expression of IKBa and IKKa (IKBa vs IKKa (log10): 0.73). Interestingly, there are correlations between soluble Gro-alpha factors and the platelet activating receptors CD62P and CD63 (e.g. GRO alpha (log10) vs CD62P−MFI (log10): 0.822; GRO alpha (log10) vs CD63−MFI (log10): 0.786), as well as RANTES and sCD62P or RANTES and Gro alpha (e.g. RANTES vs GRO alpha (log10): 0.921). Markers that consistently respond across multiple stimulation conditions (e.g., PAR-1, PAR-4, and ADP) include i) CD62P-MFI (a universal marker of platelet activation, showing strong responses across conditions), ii) AKT (a signaling marker that consistently reflects activation pathways across multiple stimuli), iii) (a chemokine involved in inflammation, displaying steady increases across various agonists) and TSLP (although not as strong as others, iv) TSLP exhibits responses under several stimulation conditions). Markers that show significant variation depending on the agonist include i) CXCL9/MIG (highly donor-dependent and shows differential expression depending on the agonist), ii) SYK(pY629/30) (specific activation seen primarily with PAR-4 stimulation), iii) NFkB1 (stimulated selectively by PAR-1 and certain inflammatory agonists), iv) Gab2 pY614 (responds variably to distinct agonists, especially ADP and PAR-4) and v) GRO-alpha (reacts differently to agonists like collagen and sCD40L). The distinction between consistent and variable markers highlights the dual role of some markers in general platelet activation (e.g., CD62P) versus those specific to certain pathways or stimuli (e.g., SYK). This insight supports the selection of specific marker panels for diagnostic or therapeutic applications tailored to different activation scenarios. Hierarchical clusters reinforce the selection of biomarker panels used in machine learning models to classify activation signatures.

This **Supplemental Figure 4** appears to illustrate the clustering of platelet responses across different stimulation conditions (e.g., unstimulated, PAR-1 agonist TRAP, PAR-4 agonist AYPGKF, ADP, collagen, fibrinogen, and sCD40L) and across different donors (labeled as d01 to d10). It likely uses methods like principal component analysis (PCA), hierarchical clustering, or heatmaps to visualize relationships in platelet activation. PAR-1 (TRAP) and PAR-4 (AYPGKF) agonists cluster separately from unstimulated conditions, indicating strong and distinct activation signatures. Responses to ADP and collagen appear more dispersed, reflecting variability in their effects across donors. While responses to certain stimuli (e.g., PAR- 1 and PAR-4) are tightly clustered, donor-specific variability is evident for other stimuli such as collagen or fibrinogen. Some donors (e.g., d03, d06) exhibit unique response patterns, suggesting individual differences in platelet reactivity. The unstimulated condition forms a distinct cluster, highlighting the baseline state of platelet activation and its clear separation from stimulated conditions. The clustering confirms that specific agonists induce distinct platelet activation profiles, supporting their potential as diagnostic or therapeutic markers. Individual differences in platelet responses suggest the importance of personalized approaches in platelet-related diagnostics and treatments and the separation of unstimulated and stimulated conditions validates the robustness of the methods used to identify activation signatures.

Heatmaps of the experiment with columns ordered by stimulation or donor showed that some variables were more stimulation-dependent, while others were more donor dependent or sample-dependent **(Supplemental Figure 5 and Supplemental Figure 6)**. The figure **(Supplemental Figure 5)** provides an analysis of platelet responses under various stimulation conditions (e.g., ADP, collagen, fibrinogen, PAR-1 and PAR-4 receptor agonists, and soluble markers like sCD40L). The data evaluates platelet activity under different types of stimulation designed to mimic clinical or experimental conditions. Stimuli include ADP, collagen, and specific receptor agonists (PAR-1 and PAR-4), which activate different signaling pathways. Measurements are taken across 10 days (d01 to d10), providing a temporal profile of platelet activity during storage. This approach captures changes due to storage lesions, activation, or degradation of functionality. The "Donor" axis highlights inter-donor differences in platelet responses, reflecting biological variability in platelet reactivity, inflammation, and immune modulation. Different stimuli elicit distinct platelet responses; i) ADP and collagen activate hemostatic pathways, as expected; ii) PAR-1 and PAR-4 agonists mimic thrombin signaling, providing insight into coagulation-linked activation and iii) Soluble markers like sCD40L suggest inflammatory or immunological roles beyond hemostasis. Variability between donors suggests the need for personalized approaches to understanding platelet biology in transfusion medicine.

The figure **(Supplemental Figure 6)** appears to illustrate the dynamics of platelet responses under various stimulation conditions across multiple days (d01 to d10).

Data is presented across 10 days (d01–d10), showing changes in platelet functionality over time due to storage effects or responses to stimuli and platelet storage lesions may influence these results, as storage impacts biochemical and functional properties. Protein phosphorylation (e.g., SYK, PLCγ2, etc.) indicates activation of signaling pathways. Markers like CD40L and CD62P (P-selectin) reflect inflammatory and activation states. Soluble mediators (e.g., RANTES, GRO-alpha) and signaling molecules show inflammatory or immune-modulatory roles. Differences in responses across donors highlight biological diversity in platelet reactivity, which may be linked to genetics, storage conditions, or donor health. Each stimulus elicits unique activation profiles, i) ADP and collagen prominently affect pathways associated with adhesion and aggregation, ii) PAR-1/4 agonists simulate coagulation-related activation with distinct phosphorylation and mediator release patterns and sCD40L may act as both a stimulus and a marker for platelet-driven inflammation. The figure showcases the complexity of platelet responses under different conditions, emphasizing the multifaceted roles of platelets in hemostasis, immunity, and inflammation. The data also underline the effects of platelet storage on their functionality, potentially impacting their efficacy in transfusions.

Then, the impact of stimulation was tested for each variable independently. We noted that only the platelet parameters AKT and sCD40L were significantly differentially expressed regardless of the agonist used for platelet stimulation (**Table 1 and Figure 2)**.

**Figure 1.**
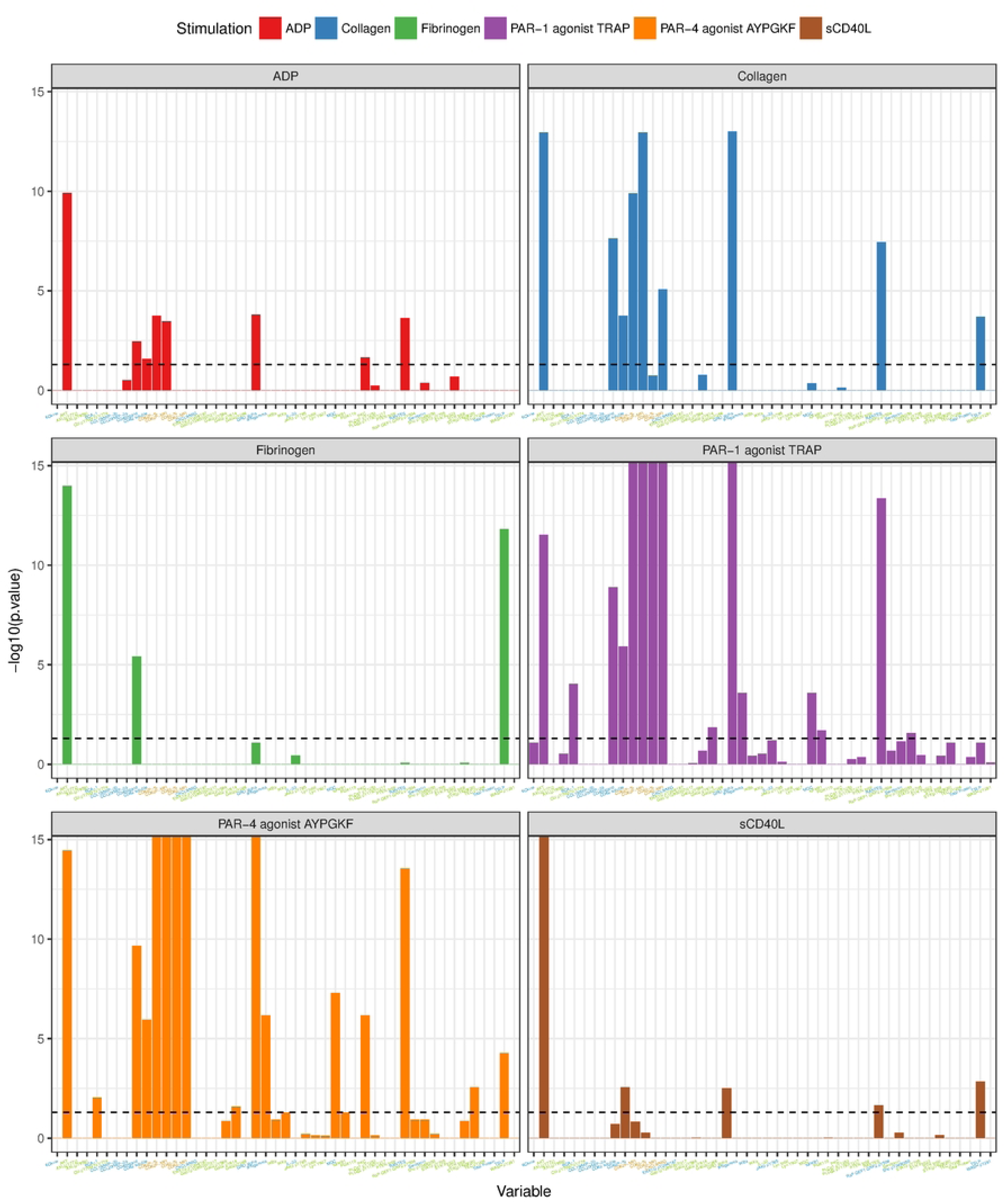
Significance of Biomarkers Across Different Biological Stimuli. This figure presents the statistical significance of various biomarkers in response to seven biological stimuli: "Unstimulated," "ADP," "Collagen," "Fibrinogen," "PAR.1.agonist. TRAP," "PAR.4.agonist.AYPGKF," and "sCD40L." The y-axis represents the negative logarithm of the p-value (-log10(p-value)), where higher values indicate greater statistical significance. Each biomarker’s contribution is analyzed, including proteins, cytokines, phosphorylation events, and other molecular markers such as CD62P, PKC, and CXCL9/MIG. The biomarkers are categorized and plotted for each condition to highlight their differential expression and relevance under various stimuli. This visualization underscores the diverse molecular pathways activated by distinct stimuli, with particular emphasis on platelet activation, immune signaling, and inflammation-related responses. The *p* values of the per-variable statistical analyses (on a - log10 scale). The dashed line represents the 0.05 threshold.

**Figure 2.**
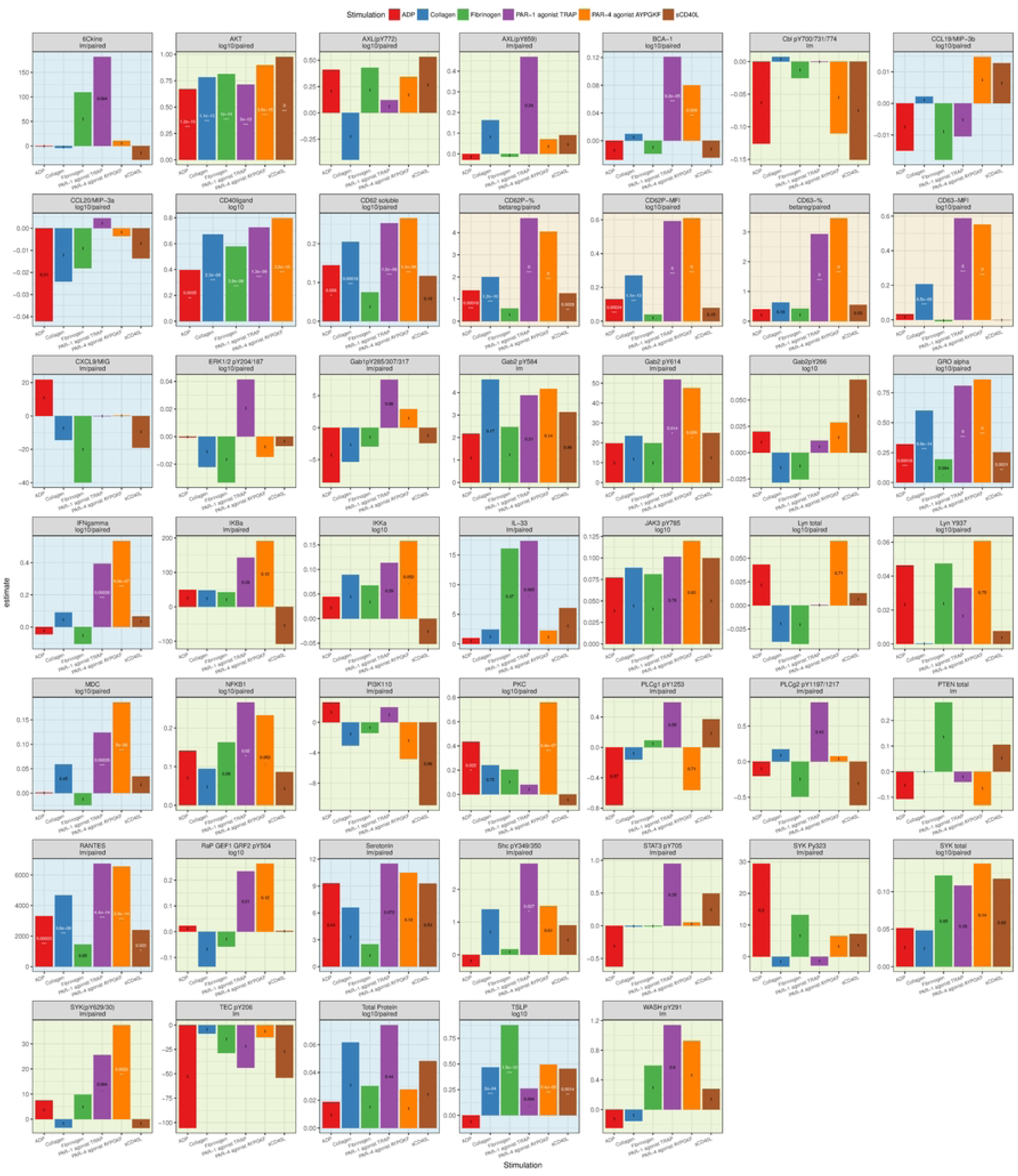
Results of the per-variable statistical analyses. The abbreviation below the variable name indicates the type of model used and stands for lm=standard linear model, log10=standard linear model after log10 transformation of the variable, and betareg=beta regression of the percentages. The symbols below the *p* values (text in the middle of the bars) indicate that the *p* value falls within the following boundaries: 0 ’***’ 0.001 ’**’ 0.01 ’*’ 0.05 ’.’ 0.1 ’ ’ 1. Additionally, *p* values below 0.05 are written in white.

**Table 1:**
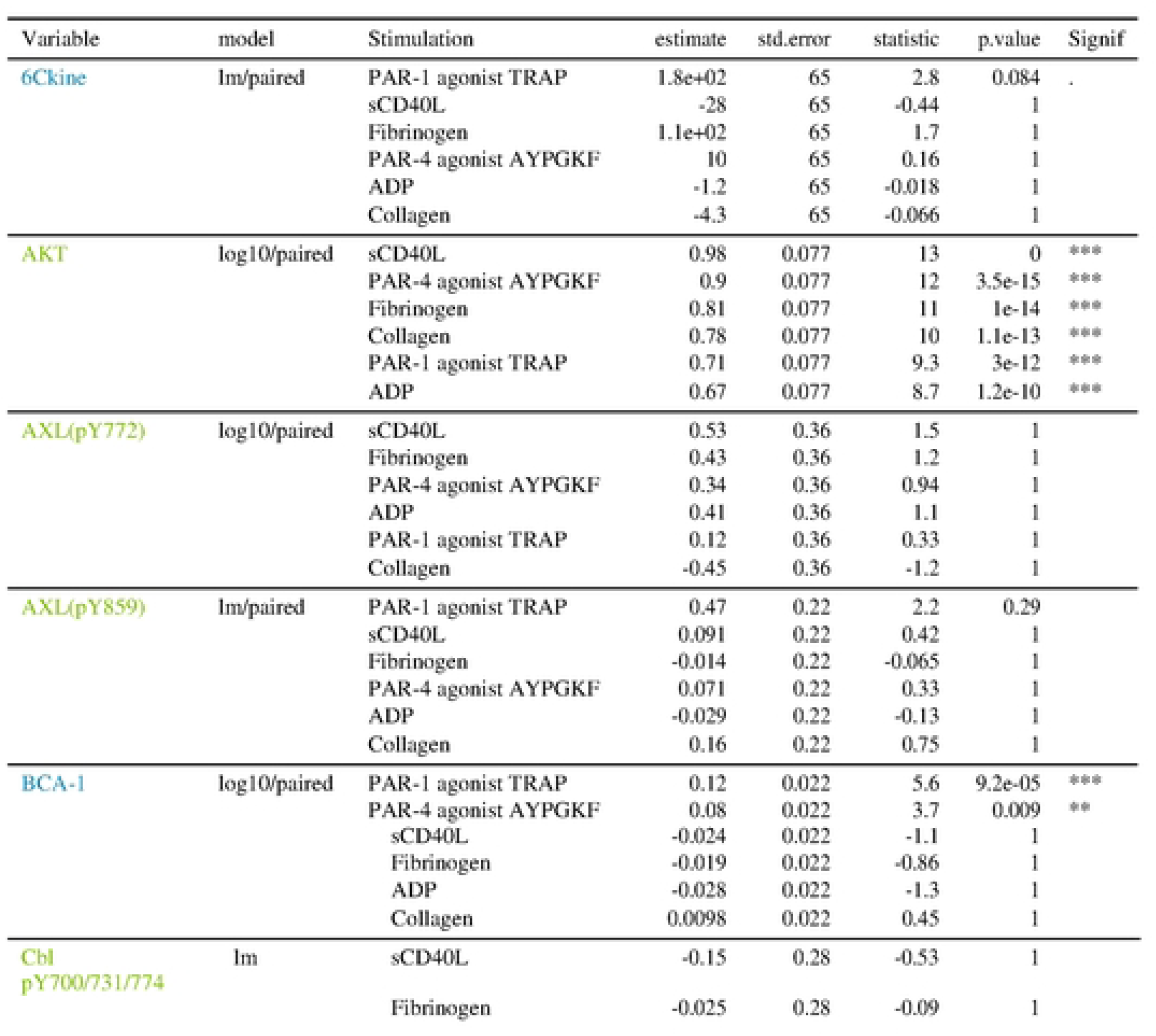
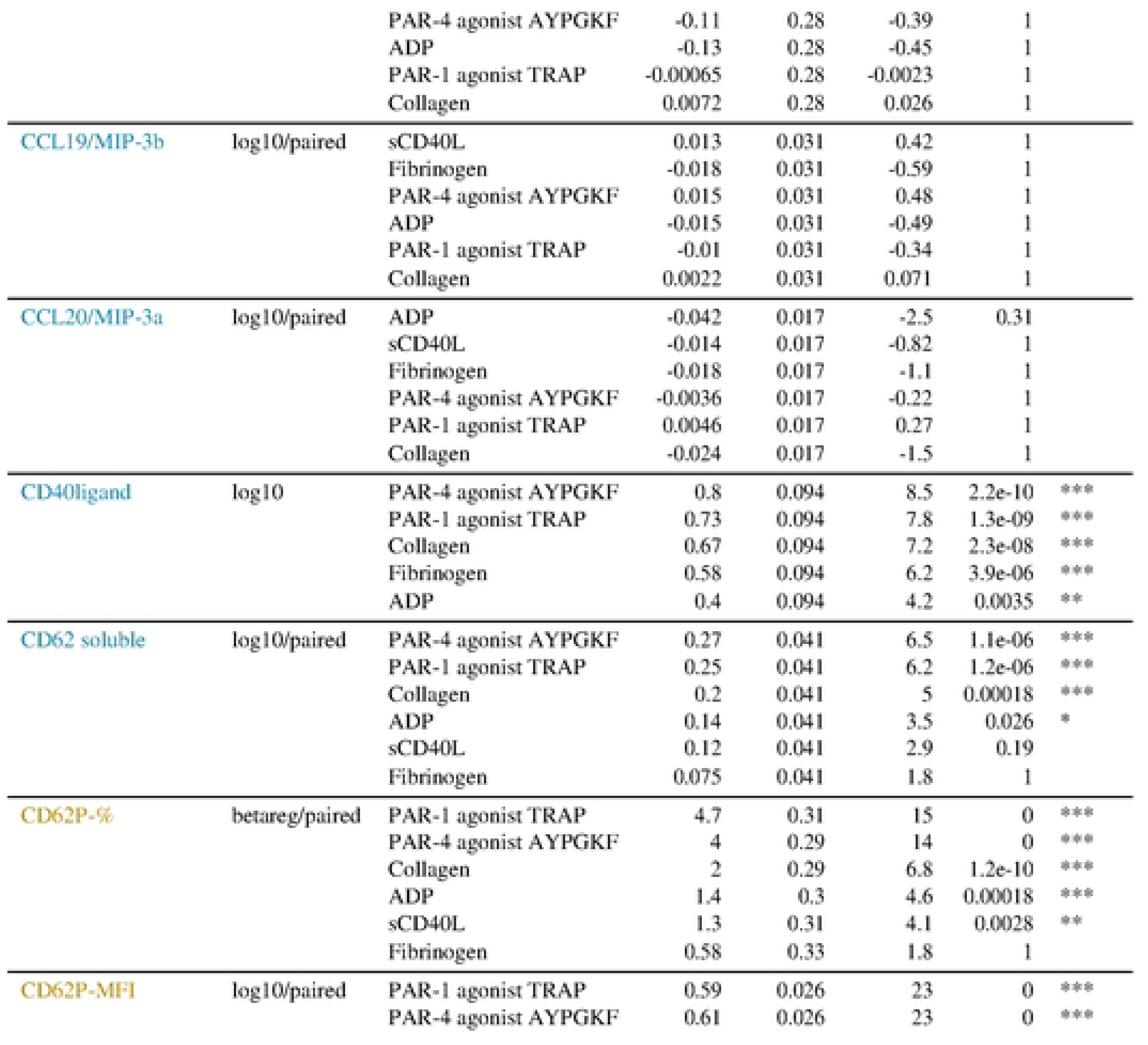
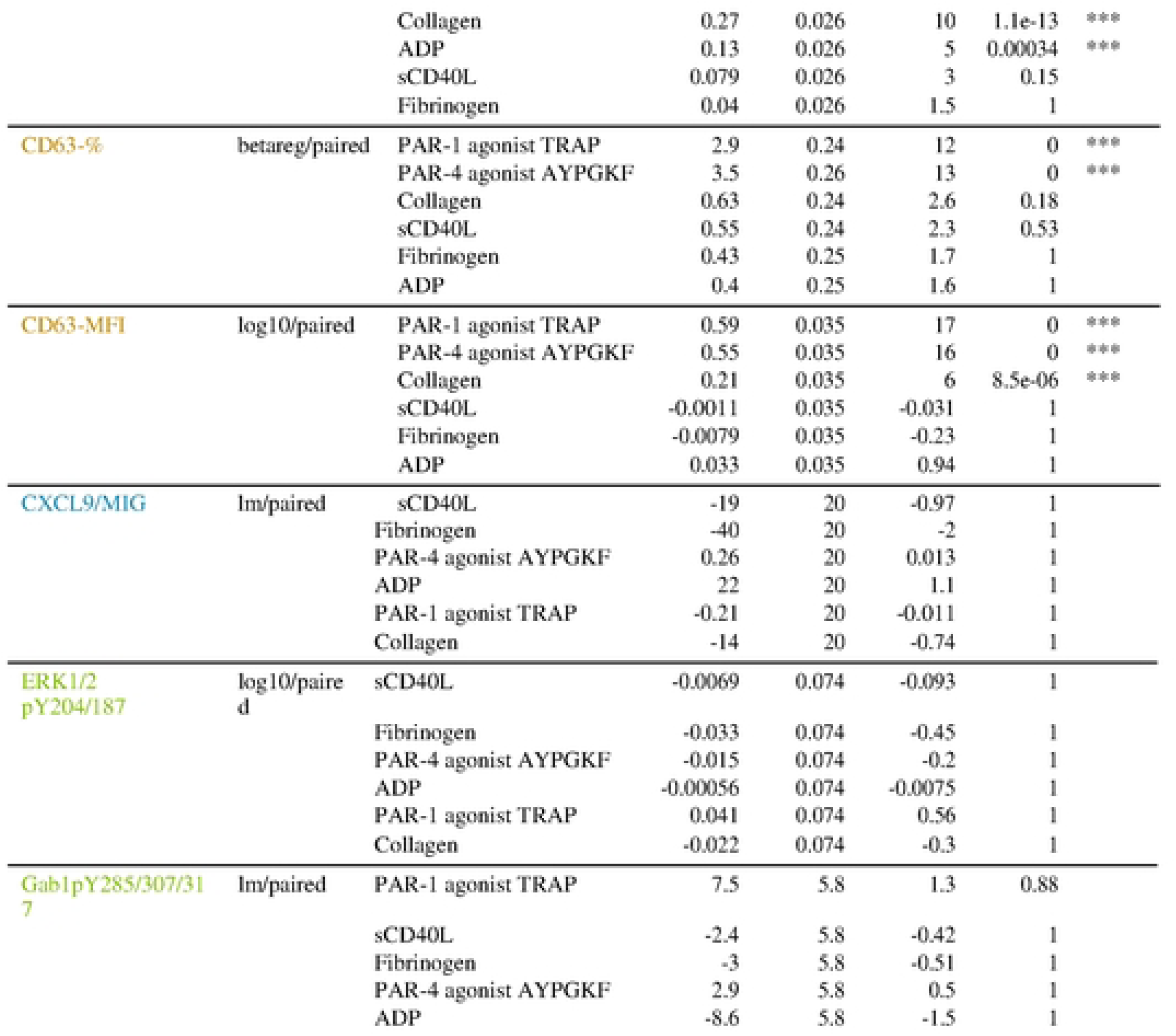
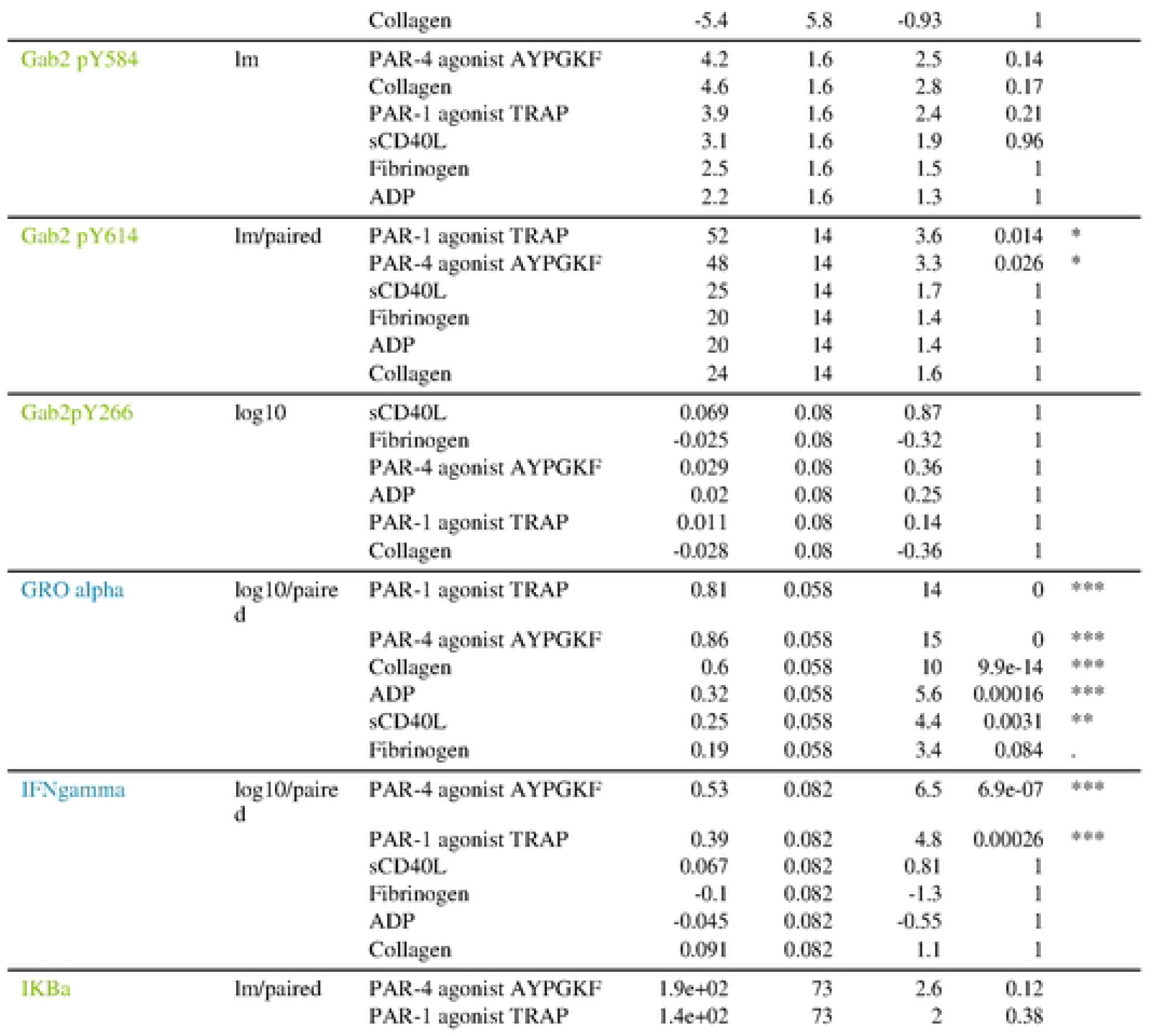
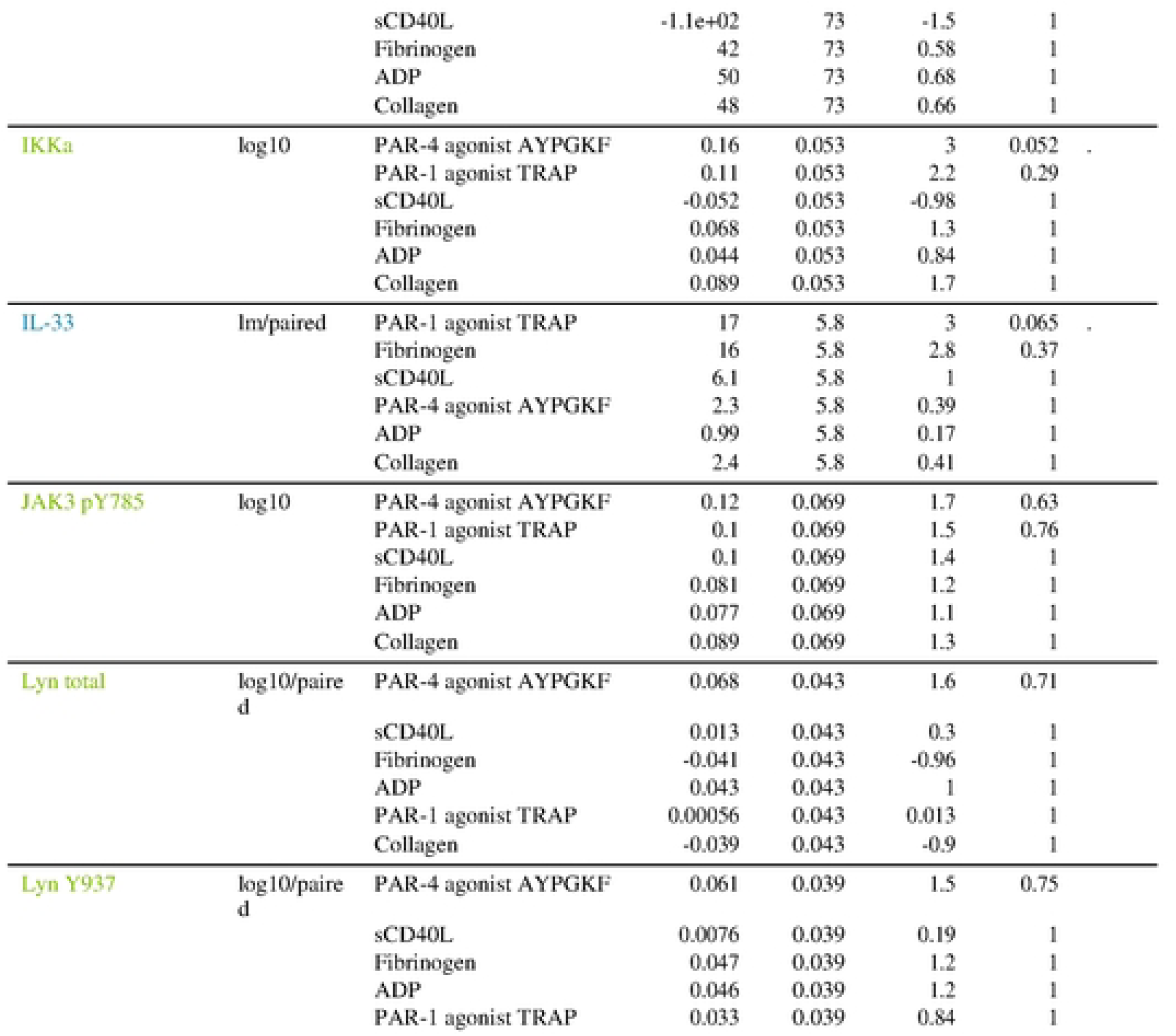
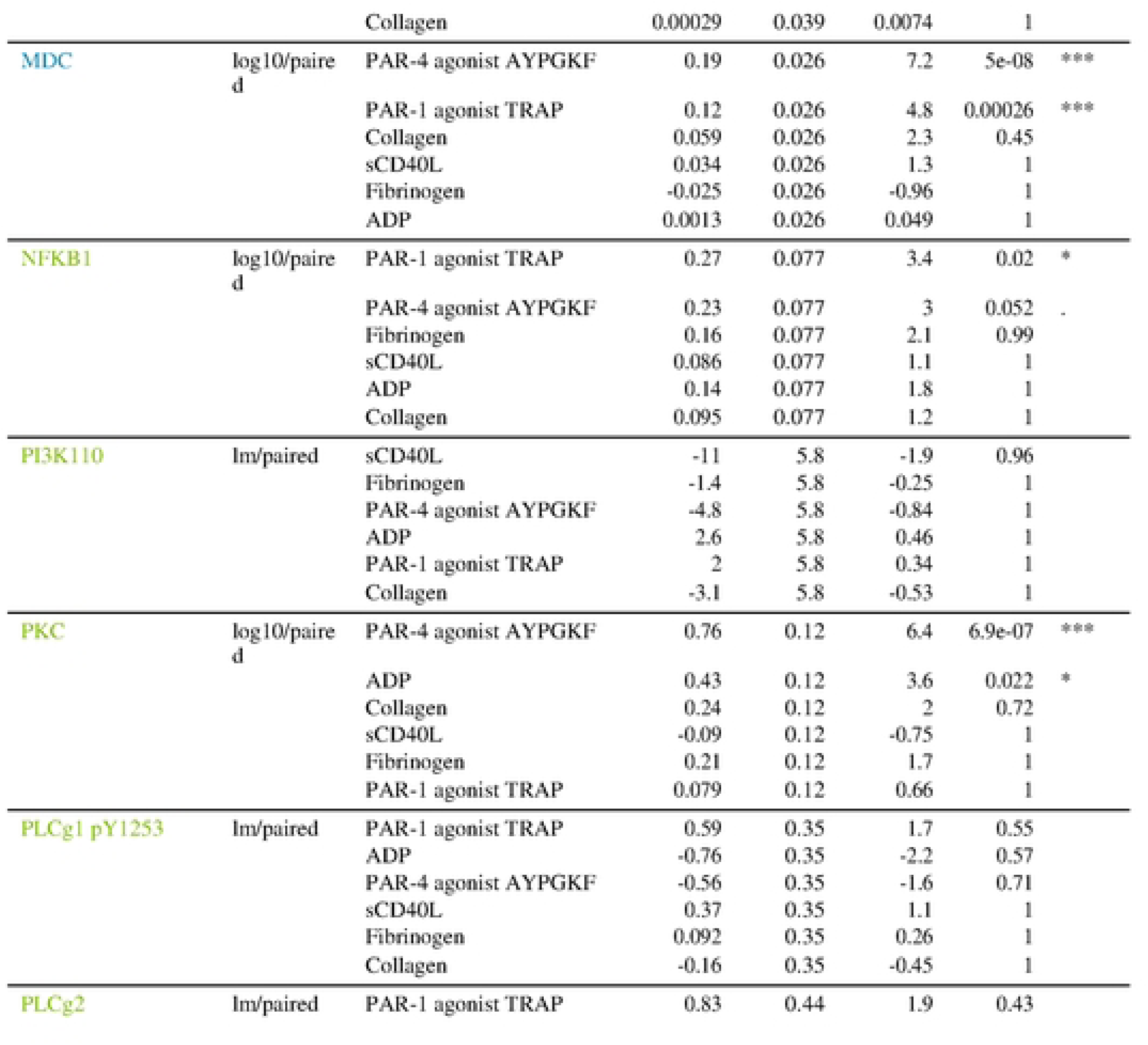
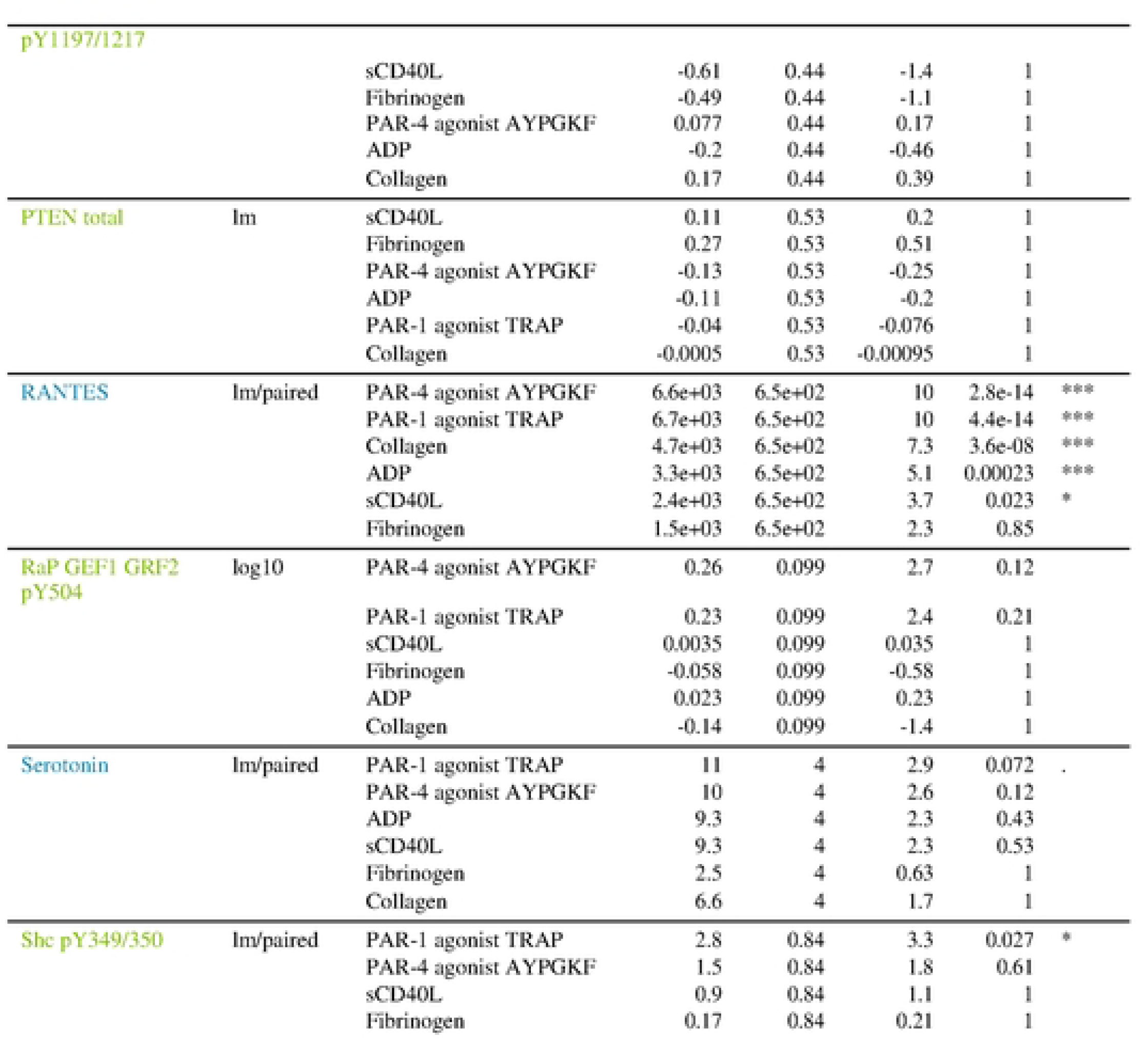
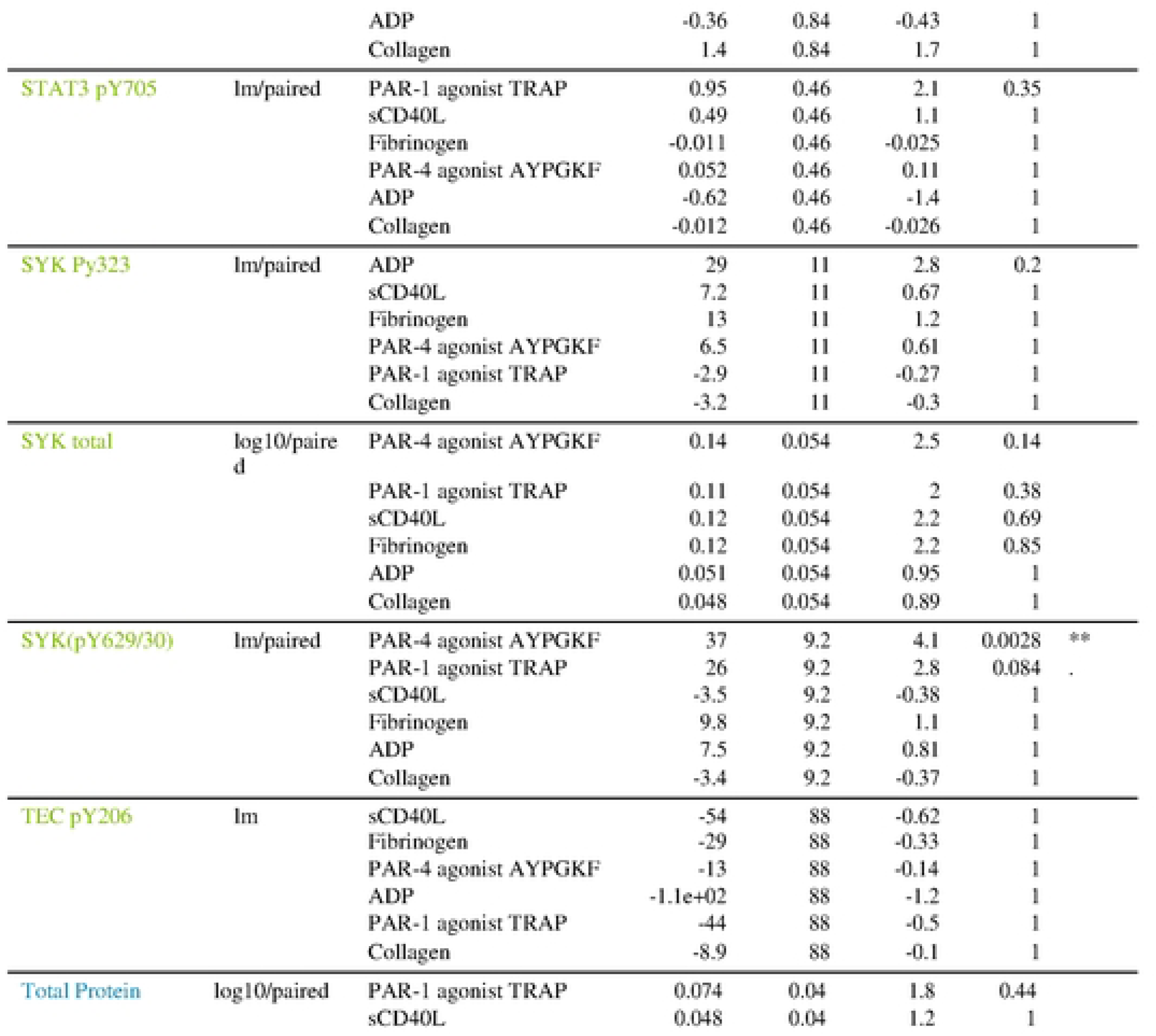
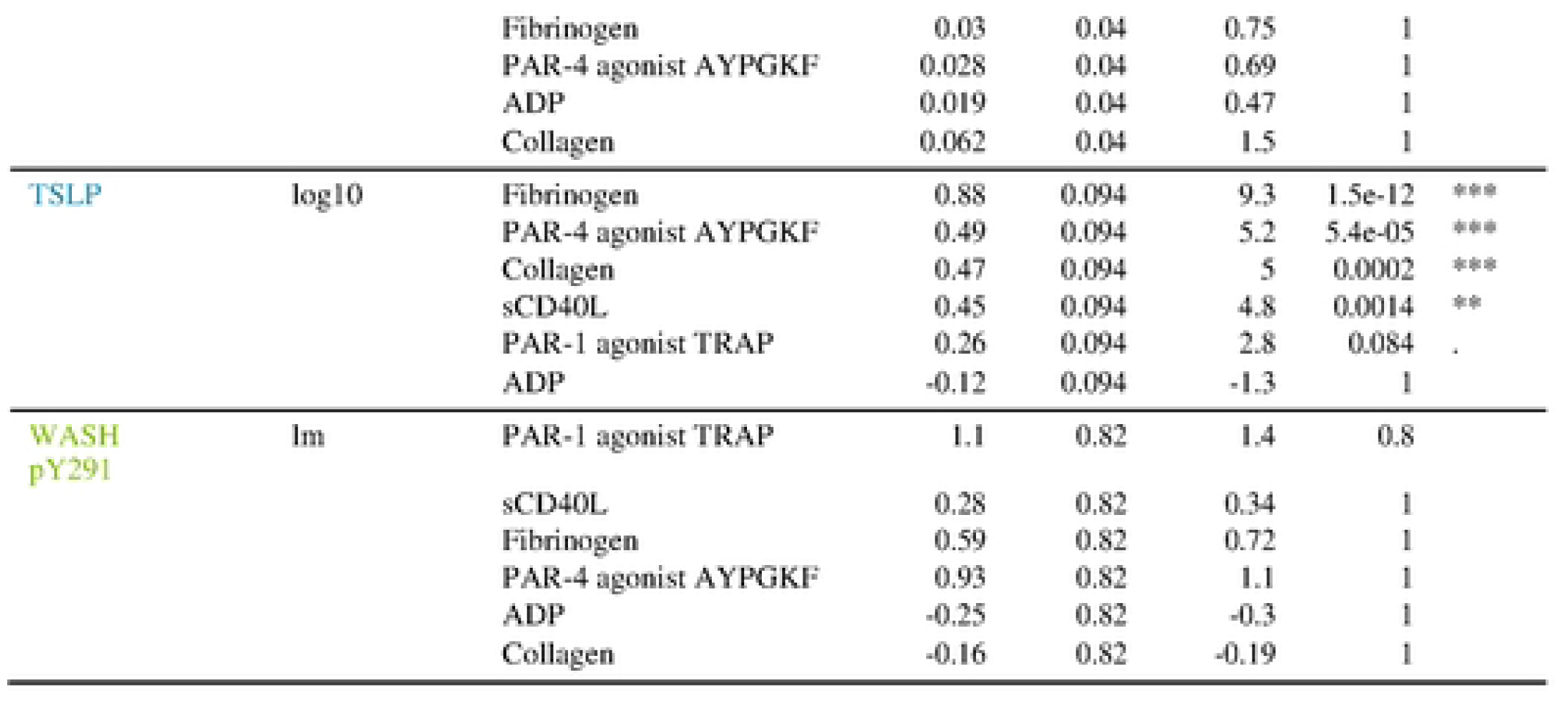
Statistical Analysis of Biomarker Responses Across Biological Stimuli. This table summarizes the statistical significance of various biomarkers under different conditions, including "Unstimulated," "ADP," "Collagen," "Fibrinogen," "PAR.1.agonist. TRAP," "PAR.4.agonist.AYPGKF," and "sCD40L." For each biomarker, the table lists the type of model used (e.g., linear regression or beta regression), the stimulation condition, the estimate, standard error, test statistic, p-value, and level of significance. For each variable, stimulations are ordered by *p* value. The “Signif” column indicates whether the *p* value falls within the following boundaries: ***p<0.001, **p<0.01, *p<0.05. Model abbreviations: lm=standard linear model, log10=standard linear model after log10 transformation of the variable, betareg=beta regression of the percentages. “Paired” was added when the donor information was included in the model.

The **Figure 2** outlines statistical analyses of responses to various biochemical stimuli across a range of factors, using metrics such as estimate values, standard errors, statistical significance, and p-values. The data provides insights into the activation or suppression of pathways under different conditions. Interestingly, IFN-gamma, BCA-1, MDC, Gab2Py614 and CD63 (% expression) were significantly and selectively differentially expressed only after platelet stimulation by the PAR-1 agonist TRAP and PAR-4 agonist AYPGKF but not by the other agonists (ADP, fibrinogen, collagen and sCD40L). The MFI of CD62P was observed for the same agonists (PAR-1 agonist TRAP and PAR-4 agonist AYPGKF) but also after collagen and ADP stimulation. TSLP was significantly released only after platelet stimulation with the PAR-4 agonist AYPGKF, fibrinogen, collagen and sCD40L. There was also an increase in GRO-alpha secretion and RANTES and CD62P expression percentages after stimulation with the PAR-1 agonist TRAP, PAR-4 agonist AYPGKF, ADP, collagen and sCD40L, while CD62P secretion and CD62P MFI were observed for the same agonists except sCD40L. Moreover, PKC was significantly increased only after stimulation with the PAR-4 agonist AYPGKF and ADP, as described previously [29]. Finally, the significant increase in SYK(Py629/30) seemed to be specifically due to platelet activation by the PAR-4 agonist AYPGKF, while that of NFkB1 and SHC(Py349/350) seemed to be specifically due to platelet activation by the PAR-1 agonist TRAP. A strong statistical association between stimulations and the activation of key molecular pathways in platelets or related cell systems. Markers such as SYK, STAT3, and Gab2 appear to be significantly activated in response to platelet agonists like TRAP, ADP, and Collagen. Some pathways show non-significant responses, indicating selective activation depending on the stimulus. The *p* values of the per-variable statistical analyses allowed analysis of parameters significantly modulated by the agonist used for platelet stimulation (**Figure 1)**. Interestingly, this analysis also revealed specific correlations between stimulation and differential expression of analytes (at a conventional p-value threshold of 0.05). We identified specific parameters among the panel of markers described above. **i)** When the ADP signaling pathway was activated, 8 parameters (AKT, PKC, CD62P (MFI and %), RANTES, sCD62P, GRO-alpha and sCD40L) were increased. ***ii)*** When the collagen signaling pathway was activated, 9 parameters (AKT, CD62P (MFI and %), CD62P (MFI), RANTES, TSLP, GRO-alpha, sCD62P and sCD40L) were increased. ***iii)*** When the fibrinogen signaling pathway was activated, 3 parameters (AKT, TSLP and sCD40L) were activated. ***iv)*** When the PAR-1 signaling pathway was activated, 13 parameters (AKT, CD62P (MFI and %), CD63 (MFI and %), RANTES, BCA-1, MDC, GRO-alpha, Gab2Py266, SHCPy349/350, NFkB1, IFN-gamma, sCD62P and sCD40L) were increased. ***v)*** When the PAR-4 signaling pathway was activated, all 16 parameters (AKT, PKC, CD62P (MFI and %), CD63 (MFI and %), TSLP, SYK629/630, Gab2Py614, GRO-alpha, RANTES, IFN-gamma, TSLP, sCD62P and sCD40L) were increased. ***vi)*** When the CD40L signaling pathway was activated, 5 parameters (AKT, CD62P (%), GRO-alpha, TSLP and RANTES) were increased (note: CD40 ligand was excluded as a readout after CD40L stimulation). Statistical modelling allowed to identify the variables significantly influenced by each stimulus. The findings highlight key pathways activated in response to platelet agonists, critical for understanding thrombosis, inflammation, and immune interactions. Highly significant markers could serve as therapeutic targets or biomarkers for disease states influenced by platelet activation.

The random forest algorithm was applied to predict the stimulus applied based on the set of variables. The confusion matrix was computed over the repeated cross-validations, with each row representing the examples in a predicted class and each column representing the examples in an actual or true class (**Supplemental Figure 7**). Variances across different conditions, especially where values are scattered (e.g., across "Fibrinogen" or "sCD40L"), could indicate challenges in prediction or variable biological responses. Most of the responses were classified into the correct groups in the diagonal line of the confusion matrix. The average accuracy was 77.6%, and the classifier had the most difficulty identifying sCD40L stimulation and differentiating between the PAR-1 and PAR-4 agonists. To further optimize the result, additional cross-validations were performed for each of the 1023 possible subsets of variables out of the set of the 10 most important variables identified in the overall classification (i.e., TSLP, CD62P-%, AKT, CD62P-MFI, CD63-MFI, CD63-%, GRO-alpha, sCD40 ligand, PKC, and RANTES) (**Supplemental Figure 8)**. The estimated accuracy of the top 10 classifiers (**Table 2)** and the confusion matrix describing the best classifiers (**Figure 3**) were determined.

**Figure 3.**
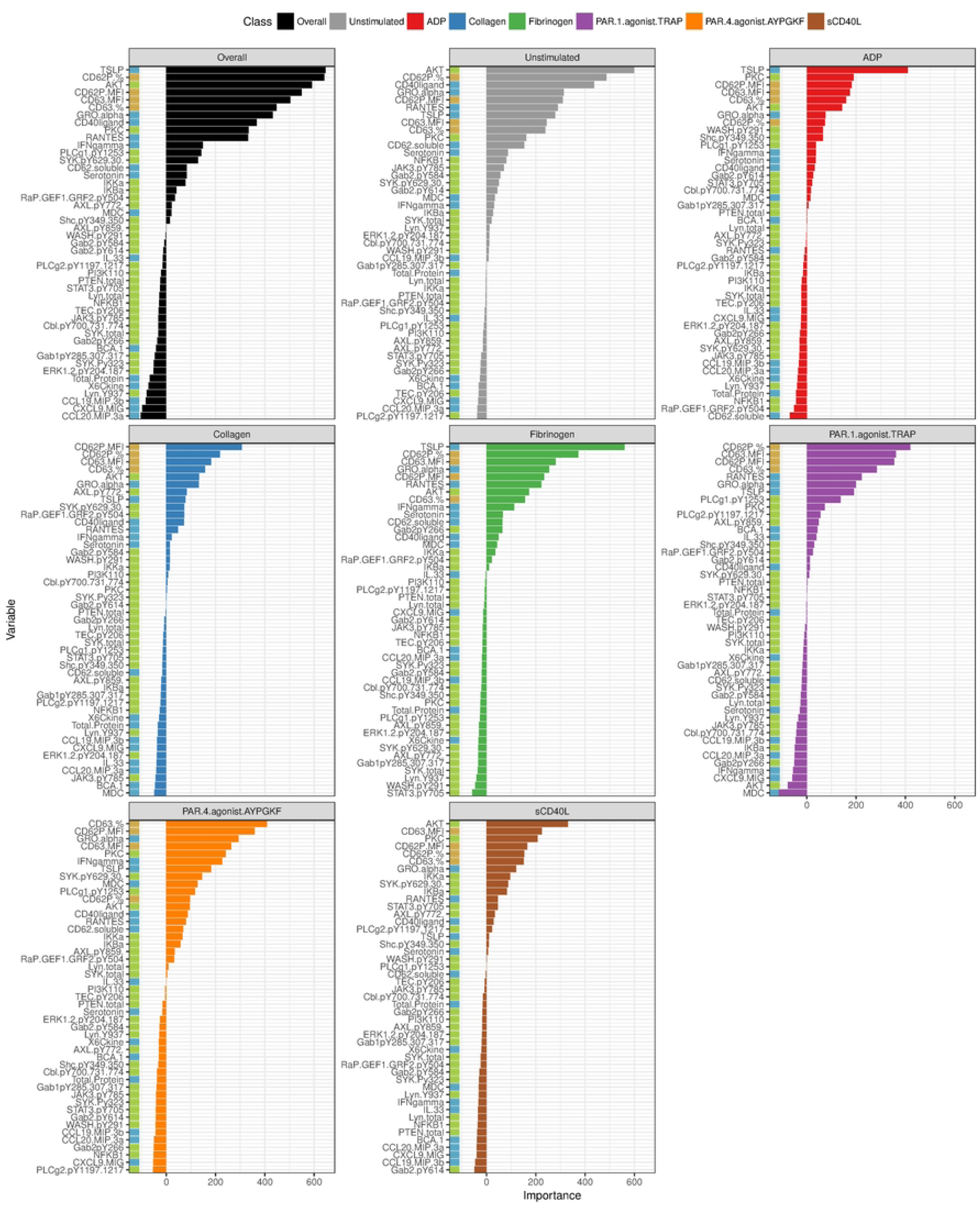
Variables by order of decreasing importance (as estimated by the “mean decrease of accuracy criterion) for the overall multiclass model and for each individual class. Variable importance as estimated by the “mean decrease in accuracy” criterion. Each panel lists, from top to bottom, the variables from most to least important for the overall (top-left panel) classification approach or for each individual class (remaining panels). The variables were arranged in order of decreasing importance for the overall multiclass model and for each individual class. The random forest was repeated with a number of trees set to 10^6^ to better estimate variable importance.

**Table 2:**
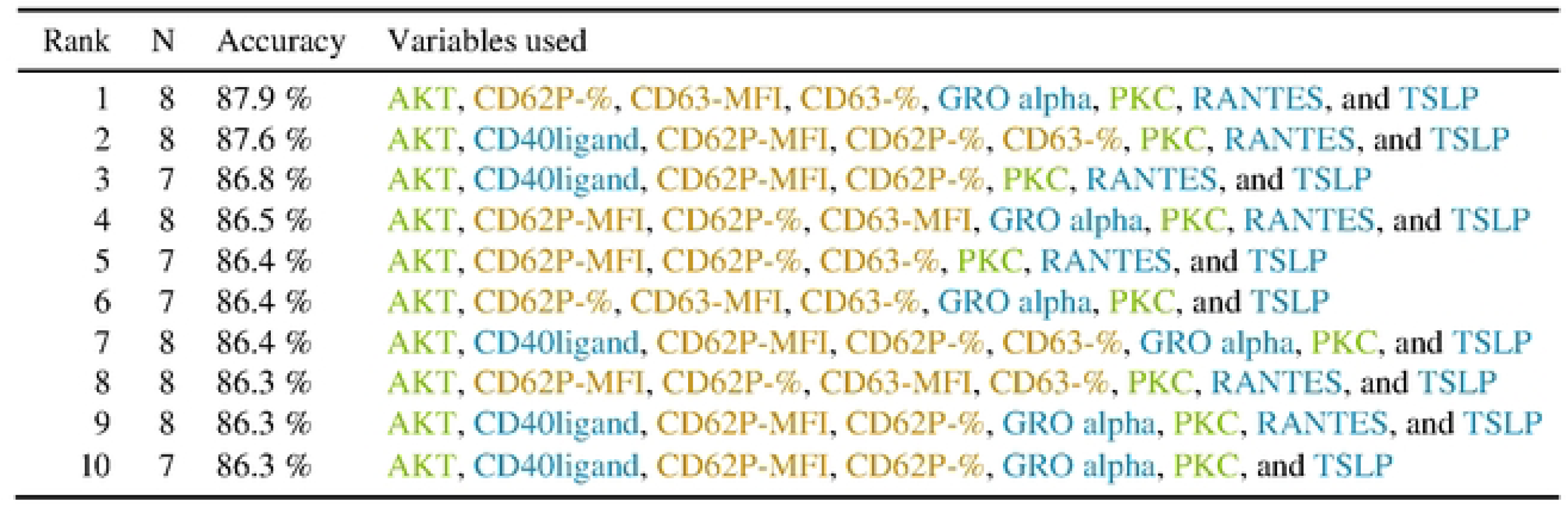
Model Accuracy and Selected Biomarkers Across Ranked Configurations. This table presents the top 10 ranked predictive models based on their accuracy and the combination of biomarkers used. Each row details the rank, the number of variables included in the model (N), the model’s accuracy (percentage), and the specific biomarkers used. The highest-ranked model achieves 87.9% accuracy using 8 variables: AKT, CD62P-%, CD63-MFI, CD63-%, GRO alpha, PKC, RANTES, and TSLP. Subsequent configurations with slight variations in biomarker selection yield similar levels of accuracy, emphasizing the robustness of core biomarkers such as CD62P, PKC, and TSLP in distinguishing biological conditions. This table highlights the importance of optimizing biomarker combinations to achieve high predictive accuracy while balancing model complexity. The colors associated with the biological parameters represent the location of the measurement: membrane activation markers (brown), soluble biological response modifiers (blue), or signaling pathway activation profiles (green).

| Rank | N | Accuracy | Variables used |
| --- | --- | --- | --- |
| 1 | 8 | 87.9 % | AKT, CD62P-%, CD63-MFI, CD63-%, GRO alpha, PKC, RANTES, and TSLP |
| 2 | 8 | 87.6 % | AKT, CD40ligand, CD62P-MFI, CD62P-%, CD63-%, PKC, RANTES, and TSLP |
| 3 | 7 | 86.8 % | AKT, CD40ligand, CD62P-MFI, CD62P-%, PKC, RANTES, and TSLP |
| 4 | 8 | 86.5 % | AKT, CD62P-MFI, CD62P-%, CD63-MFI, GRO alpha, PKC, RANTES, and TSLP |
| 5 | 7 | 86.4 % | AKT, CD62P-MFI, CD62P-%, CD63-%, PKC, RANTES, and TSLP |
| 6 | 7 | 86.4 % | AKT, CD62P-%, CD63-MFI, CD63-%, GRO alpha, PKC, and TSLP |
| 7 | 8 | 86.4 % | AKT, CD40ligand, CD62P-MFI, CD62P-%, CD63-%, GRO alpha, PKC, and TSLP |
| 8 | 8 | 86.3 % | AKT, CD62P-MFI, CD62P-%, CD63-MFI, CD63-%, PKC, RANTES, and TSLP |
| 9 | 8 | 86.3 % | AKT, CD40ligand, CD62P-MFI, CD62P-%, GRO alpha, PKC, RANTES, and TSLP |
| 10 | 7 | 86.3 % | AKT, CD40ligand, CD62P-MFI, CD62P-%, GRO alpha, PKC, and TSLP |

**Table 2** provides a ranking of models based on their predictive accuracy and the variables included. These models were likely used to predict specific biological responses or classify conditions based on platelet or immune system markers. The highest-ranked model achieves an accuracy of 87.9% using 8 variables: AKT, CD62P-%, CD63-MFI, CD63-%, GRO alpha, PKC, RANTES, and TSLP. Models ranked second and third demonstrate slightly lower accuracy (87.6% and 86.8%) with a similar selection of variables, such as CD40ligand, RANTES, and TSLP. Certain variables, including AKT, CD62P-%, CD63-%, PKC, RANTES, and TSLP, are recurrent across the top 10 models. This consistency highlights their importance in driving predictive accuracy. Models using 7 variables (e.g., Rank 3) achieve near-identical performance to those using 8 variables, suggesting that reducing the model’s complexity minimally impacts accuracy. The table demonstrates a robust machine learning framework where predictive accuracy approaches 88%. The frequent recurrence of key variables across models underlines their biological and statistical significance, suggesting their critical role in capturing the underlying mechanisms.

Condition-specific variable importance highlights distinct biomolecular pathways activated under different stimuli, emphasizing their roles in platelet function and immune response. In the "Unstimulated" condition, CD62P.% emerges as a dominant marker, consistent with baseline platelet activation profiles. Under ADP and Collagen stimulation, reliance on phosphorylation markers like PLCg2.pY1197 and signaling proteins such as PKC underscores the activation of mechanistic pathways associated with these agonists. For Fibrinogen, pivotal markers such as CD63 and CD62P reflect its direct role in platelet aggregation. In thrombin receptor pathways triggered by PAR.1.agonist. TRAP and PAR.4. agonist AYPGKF, markers like Gab2.pY584 and SYK.pY629 highlight downstream signaling specificity. The sCD40L condition, in contrast, shifts importance towards inflammatory markers like IL-33 and CD40ligand, illustrating its centrality in immune modulation. Universally important markers such as CD62P, PKC, and chemokines (CXCL9.MIG, CCL20.MIP.3a) indicate their broad involvement, while phosphorylation markers (e.g., SYK and PLCg2) show selective importance under conditions like ADP, Collagen, and PAR agonists, reflecting condition-specific activation of platelet signaling pathways (**Figure 3**).

Because platelets can be activated by multiple agonists simultaneously *in vivo*, we observed that the diagonal values in the confounding matrices (**Supplemental Figure 7** and **Supplemental Figure 8)** were not consistently 100%. However, there were some cross-coupling effects of multiple agonists on platelet activation. Finally, a multiclass classifier with an accuracy of 87.9% was obtained from a subset of 8 variables, although the number of samples was relatively small compared to the number of classes considered.

## Discussion

Our study used biomathematical approaches to identify specific differential regulation of key biological parameters of platelets in response to distinct platelet agonists. AKT, CD40 ligand, CD62P (MFI and %), PKC, RANTES and TSLP were found to be the best (limited) panel for measuring human platelet activity with an accuracy of 86.8%. Some specific correlations between specific stimuli and differential analyte expression were revealed in our analysis (i.e. CD40L signaling pathway was activated, 5 parameters (AKT, CD62P (%), GRO-alpha, TSLP and RANTES). Some of which were strongly correlated. When a stimulation effect was observed, it tended to be consistent among the various donors. Statistical models allowed the identification of variables (AKT and sCD40L) that were significantly impacted by each stimulus. Finally, although the number of samples was relatively low compared to the number of classes under consideration, a multiclass classifier with 87.9% accuracy was obtained from a subset of 10 variables. Our mathematical approach enabled us to identify a small number of platelet response markers (TSLP, CD62P-%, AKT, CD62P-MFI, CD63-MFI, CD63-%, GRO-alpha, CD40 ligand, PKC, and RANTES) that characterize different platelet activation processes (PAR-1 agonist TRAP, PAR-4 agonist AYPGKF, ADP, collagen, sCD40L, or fibrinogen). To use the lowest possible number of readouts reflecting the broadest possible platelet activation, we considered that the best (limited) panel to measure was AKT, CD40 ligand, CD62P (MFI and %), PKC, RANTES, and TSLP, which together provide an accuracy of 86.8%. Recently, different studies have presented contrasting ideas [11, 27, 30–32] about the differential secretion of various membrane, intracellular pathways or granule markers from platelets in response to different agonists. These different responses may contribute to specific processes mediated by platelets to monitor and/or regulate the roles of platelet hemostasis and innate immunity, host defense against infection and inflammation. Our data confirm this original and recent view of platelet physiology: Platelets respond differently to different stimuli.

In order to investigate the above point of view, several high technologies have been developed. Cross-talk between signaling pathways cannot be predicted by studying them in isolation, and the combinatorial complexity of multiple agonists acting together prohibits an exhaustive exploration of the complete experimental space. The study by Chatterjee *et al.* [33] utilized pairwise agonist scanning (PAS), a strategy that trains a neural network model based on measurements of cellular responses to individual and all pairwise combinations of input signals to measure calcium signaling in human platelets in EDTA-treated plasma exposed to 6 different agonists and 10 donors. The results suggested a mechanism of synergy between thrombin and thromboxane. Moreover, Recently, Zhou *et al.*[34] presented an intelligent method for classifying the aggregation of platelets by agonist type. They used four different concentrations of four agonists using platelets of four healthy human subjects for platelet activation to examine the potential influence of agonist concentrations on the ability to differentiate platelet aggregates by agonist type. The authors showed that platelet aggregates could be classified by agonist type through their morphology [34]. The present study and that by Zhou *et al.* [34] are complementary because the latter tested different agonist concentrations (n=4 for each agonist - ADP, collagen, TRAP-6, and U46619) in one healthy subject but evaluated few parameters (n=2 - platelet aggregates and flow cytometry). Our study investigated several agonists (n=7) with a single concentration, resulting in the identification of 3 agonists that were in the same concentration range as the study by Zhou *et al.* [34] with 10 healthy people, and the platelet response was measured by 47 platelet biological parameters (membrane activation markers, n=4; soluble biological response modifiers, n=15; and signaling pathway activation profiles, n=28). However, the purpose of our study and that of Zhou *et al.* [34] converge to identical results confirming platelets sense danger-signal, they regulate the resulting pro-inflammatory and pro-hemostatic functions with finely and differentially tuned signals.

More recently, the interactions of platelets and their environment beyond hemostasis, specifically in inflammation, have been a major topic of interest. It has been suggested that platelets recognize and respond to local dangers, such as infectious pathogens, at the site of wounds. It has previously been described that platelet activation is followed by secretion from at least three types of storage vesicles: dense granules, α-granules and lysosomes. There are several unresolved questions concerning platelet secretion biology, and it remains controversial how platelets organize their α-granule cargo and use their canalicular system. By quantitative immunofluorescence colocalization, Kamykowski *et al.* [35] found that pairwise comparisons of 15 angiogenic-relevant α-granule proteins displayed little, if any, pattern of functional clustering. Battinelli *et al.* [23] elegantly demonstrated that activation of human platelets with adenosine diphosphate (ADP) stimulated the release of VEGF but not endostatin, whereas thromboxane A(2) (TXA(2)) released endostatin but not VEGF. Moreover, platelet release generated upon activation with ADP promoted the migration and formation of capillary structures by HUVECs in *in vitro* angiogenesis models. Jonnalagadda *et al.* [36] presented a systematic analysis of human platelet secretion in which the release kinetics of multiple cargo molecules were examined. Detailed analysis of the release rate constants indicated that platelet release heterogeneity is kinetically based and can be defined as 3 classes of events (fast, intermediate, and slow), suggesting that platelet secretion is a stochastic process potentially controlled by several factors, such as cargo solubility, granule shape, and/or granule-plasma membrane fusion routes [36, 37].

It has been suggested that the differential release of pro- and anti-angiogenic growth factors from platelet α-granules after PAR-1 and PAR-4 stimulation may be important for the regulation of angiogenesis [38]. Our results demonstrated that the vast majority of pro- and anti-angiogenic growth factors responded similarly to PAR-1 or PAR-4 engagement. However, differences were observed in membrane CD40L expression and α-granule GRO-α and MDC secreted proteins [39]. Finally, Battinelli *et al.* used super resolution microscopy of platelet α-granules simultaneously probed for 2 different membrane proteins (VAMP-3 and VAMP-8) and multiple granular content proteins (bFGF, ENDO, TSP, VEGF) and confirmed differential packaging of protein contents into α-granules [23]. All these studies demonstrated a previously misunderstood complexity concerning the formation and secretion of platelet granules and challenge earlier conceptions of how these granules are organized within platelets and contribute to the plethora of platelet physiological functions. Our data confirm that platelet granule secretion is a stochastic process potentially controlled by several factors, as described previously by the groups of Sidney W. Whiteheart [36, 37, 40–44] and Robert Flaumenhaft [45–52].

A limitation of our study was the low classification accuracy values of some agonists in the confusion matrices due to the inability of *in vitro* experiments to elucidate all factors involved in platelet activation. Furthermore, the choice not to include, for example, TXA2 (or its analogue U46616), one of the major secondary platelet activators, or CRP, which stimulates GPVI, one of the major platelet activation pathways, is justified by the focus on the platelet inflammatory response compared to the haemostasis response. Moreover, the fibrinogen concentration in the PRP was not determined prior to stimulation. Why the addition of soluble fibrinogen to PRP stimulates platelets remains unclear. We have not yet had the opportunity to determine whether this phenomenon also occurs with the addition of a GPIIbIIIa agonist. Another hypothesis is that the fibrinogen used was not free of fibronectin. Although this was not the primary aim of the study, this observation represents a limitation of the research. In addition, the sample size of 10 individuals could be considered insufficient to establish patterns of agonist response specificity. However, the originality of this study was that all 47 biological parameters of platelets were measured for each individual.

Overall, the pleomorphic nature and heterogeneity in cargo distribution of the a-granule population presents a challenge for further study. It is challenging but is expected to be intriguing to study and elucidate the mechanisms for a further understanding of the biology of platelets. It is clear that there is a need for precise characterization of the platelet secretory pathway in order to define the criteria that control degranulation and secretion of selective mediators that promote or counteract adhesive and inflammatory responses. Finally, although the prepared PRP is very slightly contaminated with mononuclear cells, we cannot exclude the possibility that this small percentage of mononuclear cells may have influenced our results.

## Conclusions

Platelets are able to differentiate different types of danger and to generate an appropriate secretory response. This response is generally, physiologically, only what is needed to face danger or to repair a limited insult. This is indeed the normal function of platelets; moreover, part of their daily role in repairing the vascular endothelium that they patrol is to detect tears and erosions and prevent ruptures and leakage (bleeding). However, the nature or extent of secretory responses by activated platelets occasionally exceeds physiological conditions and becomes pathogenic. This suggests that there may be susceptibility factors in individuals, as well as favorable conditions, that could be relevant to disease development and/or exploited as therapeutics. This latter issue is now being considered with renewed interest because of the ease with which platelet action can be manipulated by drugs. If properly addressed, platelet activation could be converted from harmful to beneficial for the patient in a number of different infectious and inflammatory situations, such as cardiovascular disease, serious infections, autoimmune diseases, autoinflammatory diseases and cancer. Further studies are needed to understand i) the process of platelet activation and release products, in the clinical context in which platelet release products have been measured as well as the potential clinical and/or diagnostic relevance and ii) the mechanisms of these fascinating cells. Finally, we plan to perform studies in platelets treated with selective inhibitors of identified platelet activation pathways or in patients with specific defects in these pathways.

## Methods

### Preparation of blood samples and platelet stimulation

The average age of a combined group of men (n = 5) and women (n = 5) is 35 years [26–43]. Peripheral blood samples from healthy subjects (n=10) were collected in endotoxin-free tubes with 3.2% sodium citrate (Vacutainer®, Becton Dickinson, San Jose, California) [39]. The blood samples were centrifuged at 282 × *g* for 10 min at RT to obtain PRP. The platelet count was adjusted in the PRP and was consistent across all groups for all investigations. The WBC count is therefore significantly reduced (often below 2.2×10³/mm³). PRP was incubated (30 min, RT), with moderate agitation (100 rpm), with (or without) the PAR-1 agonist TRAP (5 µg/ml), PAR-4 agonist AYPGKF (0.2 mM), ADP (10 µg/ml), fibrinogen (100 μg/mL) (Sigma Aldrich, St. Louis, USA), collagen (1 mg/ml) (Nycomed, Munich, Germany) or sCD40L (100 µg/ml) (Enzo Life Sciences, Villeurbanne, France). Platelets were processed immediately for Flow cytometry assay (membrane activation markers). The PRP was centrifuged at 402g for 10 minutes at room temperature. The supernatant, aliquots were frozen at -80°C (releasome).

The platelet pellet was washed and intracellular proteins were extracted using a MILLIPLEX MAP EpiQuant Sample Preparation kit at 3 × 10^7^ platelets/ml of lysis buffer, according to the manufacturer’s instructions and at -80°C [39].

## Methods of measuring platelet activity

### Flow cytometry assay

PRP samples with or without stimulation were incubated at room temperature with antibodies against CD41a (clone HIP8, BD Biosciences, Le Pont de Claix, France), CD62P (clone AK-4, BD Biosciences) and CD63 (clone H5C6, BD Biosciences) for 30 min at RT. Cells were fixed with ThromboFix reagent (Beckman Coulter, Villepinte, France) for 1 h at room temperature according to the vendor’s instruction and washed once with phosphate-buffered saline (1×) prior to analysis with a FACSVantage SE device equipped with CellQuestS-Pro software (BD Biosciences, Le Pont de Claix, France).

### Enzyme-linked immunosorbent assay (ELISA)/Luminex

The α-granule protein content of platelet supernatants was quantified using Luminex® technology with the following panel of human cytokine/chemokine magnetic beads: HCYTOMAG-60K, HCYP2MAG-62K, and HCYP3MAG-63K (Millipore, Molsheim, France). A canonical δ granule molecule, serotonin (outside of the panel described above), was quantified by ELISA (IBL International, Hamburg, Germany). Absorbance at 450 nm (or 405 nm for serotonin) was measured using an ELISA plate reader (Magellan Software, Sunrise TM, Tecan Group Ltd., Lyon, France). The results were normalized to 10^9^ platelets/ml.

For each condition, intracellular proteins from the platelets were extracted using a MILLIPLEX MAP EpiQuant Sample Preparation kit at 3 × 10^7^ platelets/ml of lysis buffer, according to the manufacturer’s instructions. The quantification of phosphorylated proteins in the platelet pellet, was performed using MILLIPLEX MAP EpiQuant technology with the following five panels: MPEQMAG-100K, 102K, 103K, 104K and 110K (Millipore; see **Supplemental Figure 1**).

In some figures and tables, the color code is always the same. The colors associated with the biological parameters represent the location of the measurement: membrane activation markers (brown), soluble biological response modifiers (blue), or signaling pathway activation profiles (green).

### Statistical and computational analysis

Variables expressed as percentages were logit-transformed. To improve normality, and based on the log-likelihood of the Box-Cox power transform (**Supplemental Figure 1** and **Supplemental Table 1)**, some of the remaining variables were log_10_ transformed. To allow this computation, values at zero were replaced with half of the minimal observed value for the given variable.

Since the experimental procedure did not allow the measurement of the CD40 ligand in the sCD40L-stimulation condition, the corresponding values were imputed from the other variables using the Non-linear Iterative Partial Least Squares (NIPALS) algorithm [53].

Percentage variables were analyzed using beta regression. Variables that were more normally distributed after a log_10_ transformation were analyzed using ordinary least squares models on that scale. The remaining variables were analyzed using ordinary least squares models on their original scale. The formulas used were the same for the three types of variables: either “value ∼ stimulation + donor” (paired model) or “value ∼ stimulation” (unpaired model). The choice was made based on the value of the AIC for each model. These models were used to estimate the effect of each stimulation compared to the outcome in the unstimulated condition. The *p* values were first corrected for multiple testing using Dunnett’s method for each variable and then corrected for each stimulation by the Benjamini and Hochberg method (FDR).

Random forests were applied to the imputed dataset. The number of trees was set to 10^4^, and the number of variables randomly sampled as candidates at each node was fixed to 6 (default value for classification given the number of variables). Model performance was estimated by repeated (100 times) stratified 5-fold cross-validations. Variable importance was estimated by the “mean decrease in accuracy” criterion using a random forest with the number of trees set to 10^6^.

### Ethics approval and consent to participate

This study complies with all relevant ethical regulations. Fresh blood from healthy donors was obtained from Etablissement Français du Sang (EFS), eligible for blood donation, upon approval of the project by Ministry of Higher Education and Research (authorization number CODECOH DC-2019-3803 & AC-2020-3959). and the French data protection law (Law n°78-17 on 06/01/1978 and Décret n°2019-536 on 29/05/2019). Informed consent was obtained from all subjects prior to the blood donation from the Etablissement Français du Sang. The study was approved by the national review board for biomedical research in March 2017 (Comité de Protection des Personnes Sud-Est I, Saint-Etienne -, France; ID RCB Number: 2014-A00405-42), in agreement with the General Data Protection Regulation (Regulation (EU) 2016/679 and Directive 95/46/EC) and the French data protection law (Law n°78-17 on 06/01/1978 and Décret n°2019-536 on 29/05/2019).

## Availability of data and materials

The datasets generated or analysed during the current study are not publicly available but are available from the corresponding author on reasonable request.

## Competing interests

All authors declare no competing financial interests.

## Funding

This work was supported by grants from the French National Blood Establishment – EFS, the Association “Les Amis de Rémi” Savigneux, France and the French Agence Nationale de la Recherche (ANR), under grant (ANR-22-CE17-0063) (project HEASY_PLAT); FC, KAN, OG, and HHC have been named as inventors in the patent application “METHOD FOR DETECTING INFLAMMATION-RELATED PLATELET ACTIVATION” (European Patent Office - Publication/Patent Number: EP4147055A1). None of the inventors received royalties from the patent application.

## Authors’ contributions

FC, KAN, HHC and OG developed the study hypothesis, designed the protocol, trained personnel, and contributed to the writing of the manuscript. SdB and JN performed data analysis and interpretation and contributed to the writing of the manuscript. FC, KAN, HHC, MH, ACD, SA and OG wrote the manuscript. KAN, FC, CAA, MAE and AP collected samples and performed the experiments.

## Acknowledgements

We would like to thank the medical staff and personnel of the Etablissement Français du Sang Auvergne-Rhône-Alpes, Saint-Etienne, France for technical support throughout our studies.

---

**Supplemental Table 1** – 𝜆 parameter selected for the Box-Cox transformation of each variable. The colors associated with the biological parameters represent the location of the measurement: membrane activation markers (brown), soluble biological response modifiers (blue), or signaling pathway activation profiles (green).

**Supplemental Table 2** – Summary statistics of the transformed variables. The colors associated with the biological parameters represent the location of the measurement: membrane activation markers (brown), soluble biological response modifiers (blue), or signaling pathway activation profiles (green).

**Supplemental Figure 1** – Log-likelihood profile for each variable for various values of the 𝜆 parameter of the Box-Cox power transformation. The vertical dotted lines represent the location of the highest likelihood and its 95% confidence interval. The colors associated with the biological parameters represent the location of the measurement: membrane activation markers (brown), soluble biological response modifiers (blue), or signaling pathway activation profiles (green).

**Supplemental Figure 2** - Distribution of the variables after log or logit transformation. Each color represents a stimulation condition. The y-axis corresponds to the count and x-axis to the value (quantification of each marker).

**Supplemental Figure 3** - Pairwise comparison of the centered and scaled variables after transformation. For each pair of variables, the values are compared in all the samples (top right) and the corresponding correlation coefficient is reported (bottom left). Perfectly correlated variables would draw a diagonal line and would have a correlation coefficient of 1 (bright green square).

**Supplemental Figure 4 -** Hierarchical clustering of samples achieved using the Euclidean distances between samples and Ward’s agglomerative method. The percentage above each cluster divide represents the stability of the grouping as estimated by multiscale bootstrap resampling.

**Supplemental Figure 5 –** Heatmaps of the stimulation experiment results (centered and scaled variables were clustered using the “complete” agglomerative method). Colors represent the intensity of the signal for each variable from low (green) to medium (yellow) to high (red). Ordered by stimulation.

**Supplemental Figure 6 –** Heatmaps of the stimulation experiment results (centered and scaled variables were clustered using the “complete” agglomerative method). Colors represent the intensity of the signal for each variable from low (green) to medium (yellow) to high (red). Ordered by donor.

**Supplemental Figure 7 - Comparison of Predicted and Reference Conditions Across Biological Stimuli**. The classifier confusion matrix illustrates the concordance between predicted and reference conditions for various biological stimuli, including "Unstimulated," "ADP," "Collagen," "Fibrinogen," "PAR.1.agonist. TRAP," "PAR.4.agonist.AYPGKF," and "sCD40L." The rows represent reference conditions, while the columns correspond to predictions made by the model. The color intensity and numerical values within each cell indicate the percentage frequency of matches between predictions and references. Diagonal elements with high frequencies reflect accurate predictions, whereas off-diagonal elements represent misclassifications or overlaps between conditions. Key observations include near- perfect prediction accuracy for "Unstimulated" and "Fibrinogen," moderate accuracy for "Collagen" and "ADP," and lower agreement for "sCD40L," indicating challenges in differentiating this condition. This figure highlights the model’s performance and its ability to discriminate between activation pathways and inflammatory responses. Each square shows the percentages of the predicted class (X-axis) for each of the held-out observations (of the class shown on the Y-axis) during cross-validations. Random forests were applied to the imputed dataset. The confusion matrix is computed over the repeated cross-validations (entries are the average percentages of the held-out observations in each cell).

**Supplemental Figure 8 - Classifier confusion matrix displays a heatmap comparing the predicted versus reference categories for different biological conditions:** The random forest was repeated with a number of trees set to 10^6^ to better estimate variable importance. To further optimize the result, new repeated cross-validations were performed for each of the 1023 possible subsets of variables out of the set of the 10 most important variables identified previously in the overall classification problem (*i.e*.: TSLP, CD62P-%, AKT, CD62P-MFI, CD63- MFI, CD63-%, GRO alpha, CD40ligand, PKC, and RANTES). Confusion matrix of the best classifier. Each square shows the percentages of the predicted class (X-axis) for each of the held-out observations (of the class shown on the Y-axis) during cross-validations. Rows represent the reference (ground truth) conditions, and columns indicate the model’s predictions. The color intensity within each cell corresponds to the percentage frequency of matches between predicted and reference values, ranging from 0 to 100%. High frequencies along the diagonal indicate accurate predictions, such as near-perfect alignment for "Unstimulated" and "Fibrinogen," whereas off-diagonal elements represent instances of misclassification, including moderate accuracy for "Collagen" and lower agreement for "sCD40L." This visualization highlights the model’s performance in differentiating platelet activation pathways and inflammatory markers, with particular strengths in thrombin receptor agonists and baseline activation, and areas requiring improvement for immune-mediated responses like "sCD40L."

## Notes

### Competing Interest Statement

The authors have declared no competing interest.

